# A dual transcript-discovery approach to improve the delimitation of gene features from RNA-seq data in the chicken model

**DOI:** 10.1101/156406

**Authors:** Mickael Orgeur, Marvin Martens, Stefan T. Börno, Bernd Timmermann, Delphine Duprez, Sigmar Stricker

## Abstract

The sequence of the chicken genome, like several other draft genome sequences, is presently not fully covered. Gaps, contigs assigned with low confidence and uncharacterized chromosomes result in gene fragmentation and imprecise gene annotation. Transcript abundance estimation from RNA sequencing (RNA-seq) data relies on read quality, library complexity and expression normalization. In addition, the quality of the genome sequence used to map sequencing reads and the gene annotation that defines gene features must also be taken into account. Partially covered genome sequence causes the loss of sequencing reads from the mapping step, while an inaccurate definition of gene features induces imprecise read counts from the assignment step. Both steps can significantly bias interpretation of RNA-seq data. Here, we describe a dual transcript-discovery approach combining a genome-guided gene prediction and a *de novo* transcriptome assembly. This dual approach enabled us to increase the assignment rate of RNA-seq data by nearly 20% as compared to when using only the chicken reference annotation, contributing therefore to a more accurate estimation of transcript abundance. More generally, this strategy could be applied to any organism with partial genome sequence and/or lacking a manually-curated reference annotation in order to improve the accuracy of gene expression studies.

## Introduction

Since its first release in 2004 and despite significant improvements over the last past decade, the *Gallus gallus* genome is presently incomplete and highly fragmented [1]. The chicken karyotype is composed of 38 autosomal chromosomes (1-38) and 2 additional sex chromosomes (W, Z) [2]. Out of these autosomal chromosomes, 10 are macrochromosomes (1-10), with lengths similar to those in mammals, and 28 are microchromosomes (11-38), with lengths ranging from 2 to 25 Mb [1]. Chicken microchromosomes display a high recombination rate, contain an elevated number of repetitive elements and are GC-rich, which induces significant bias and sequencing errors when using high-throughput technologies [3,4]. In addition, microchromosomes are gene dense and enriched in CpG islands, which is the result of short intronic sequences [5,6]. The fourth version of the *Gallus gallus* genome (galGal4) released in November 2011 has not fully overcome these issues. Out of the 40 chromosomes, 31 are sequenced (1-28, 32, W, Z) and contain more than 9,000 gaps, while 9 chromosomes remain missing (29-31, 33-38). The genome is also composed of about 16,000 additional contigs that are not assigned to any chromosome or assigned with low confidence. In total, the galGal4 genome sequence has a size of 1.05 Gb.

RNA sequencing (RNA-seq) data processing and results are highly dependent on the quality of the genome sequence and the associated gene annotation model. Read mapping is one of the critical steps that will further influence sample normalization, gene expression quantification and the identification of relevant genes. Gene expression profiles rely on the alignment of RNA-seq reads along the available reference genome or transcriptome, followed by their assignment to gene features. An incomplete genome sequence coupled with an inaccurate definition of gene features induce a bias in the gene expression quantification and transcript abundance estimation [7,8]. Whole transcriptome sequencing offers valuable resources to detect novel genes and transcripts as well as to identify alternative splicing variants [9,10]. Depending on the context, two main strategies are widely used to analyse RNA-seq data [11]. One approach consists of the mapping of reads along the reference genome followed by gene prediction [8,12,13]. This method can be coupled with an existing reference annotation in order to detect new transcripts with respect to the provided gene annotation model [14]. The second approach aims at reconstructing the whole transcriptome independently of the reference genome [15-17]. This method is particularly suitable to study models with partial or missing genome sequence. The choice between these approaches greatly depends on the biological question and whether a reference genome is available [18].

When analysing RNA-seq data obtained from chick embryonic limb cell cultures (so-called micromass cultures) by using the galGal4 reference genome and annotation, we observed that only 62.2% of sequencing read pairs were assigned to gene features, while 86.7% of the read pairs were mapped against the genome sequence. By comparison with the human genome, which has been nearly completely sequenced and accurately annotated, a similar analysis of RNA-seq data obtained from human blood samples depicted an assignment rate to gene features of 81.8% with a mapping rate of 92.3% [19]. We hypothesized that information was lost during the analysis of chick RNA-seq data: (i) at the mapping step, either due to low-quality sequencing reads, or due to missing genome sequence; and (ii) at the read assignment to gene features, which can be due to missing or partially annotated transcripts. To address both issues, we performed a dual transcript-discovery approach by means of genome-guided gene prediction and *de novo* transcriptome assembly. The approach described here enabled us to increase the assignment rate of RNA-seq data by nearly 20% as compared to when using the chicken reference annotation, thus contributing to a more robust quantification of gene expression profiles.

## Results

We performed RNA-seq of two independent biological replicates of chick micromass cultures infected for 5 days with empty RCAS-BP(A) replication-competent retroviral particles. 61.3 and 70.3 million of strand-specific read pairs were generated and mapped against the galGal4 version of the chicken genome (Table 1). Read assignment was performed by using a gene annotation model composed of 17,318 genes resulting from the combination of both UCSC and Ensembl reference annotations that were available at the time of analysis. Surprisingly, while 86.7% of read pairs were mapped against the chicken genome, only 62.2% of read pairs were assigned to gene features (Table 1). Therefore, 28.3% of mapped read pairs were not counted, including 93.7% of these read pairs that were not overlapping with any gene feature (Table 1). Close investigation of these unassigned read pairs highlighted genes that seemed to be absent or partially covered by the UCSC and Ensembl reference annotations (Fig. 1a, b), as well as transcripts with missing or partial exon features (Fig. 1c).

**Table 1.**
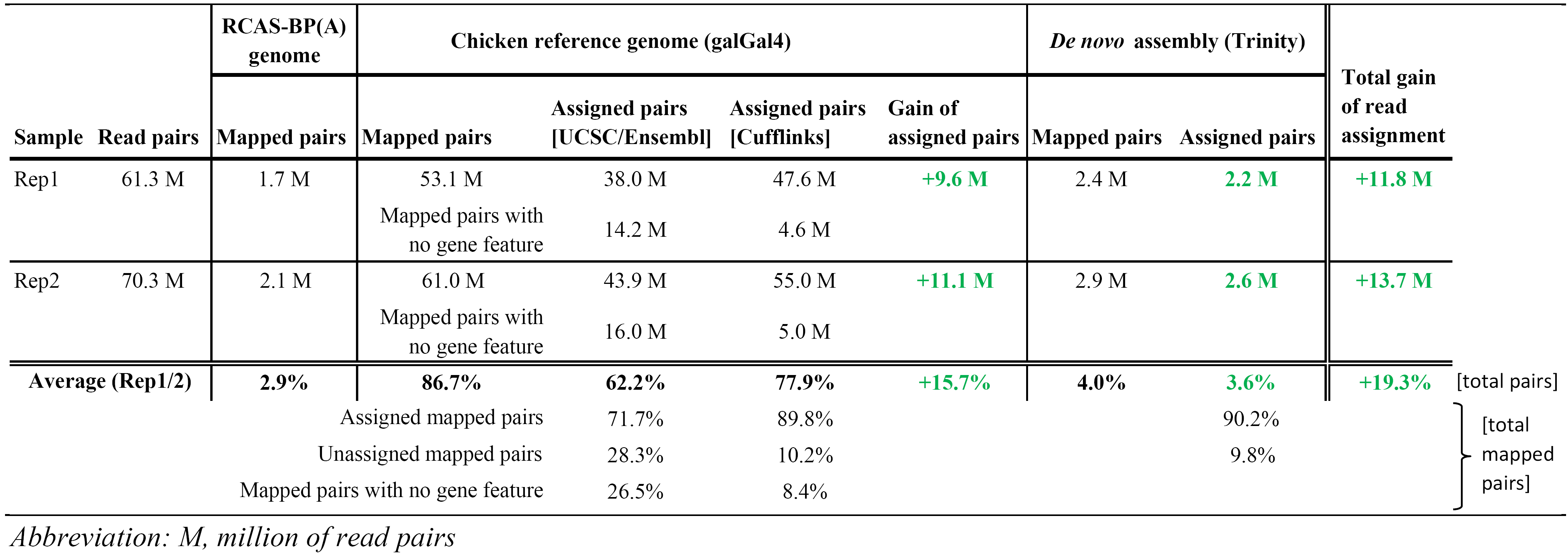
Statistics of RNA-sequencing read pair assignment

**Fig. 1.**
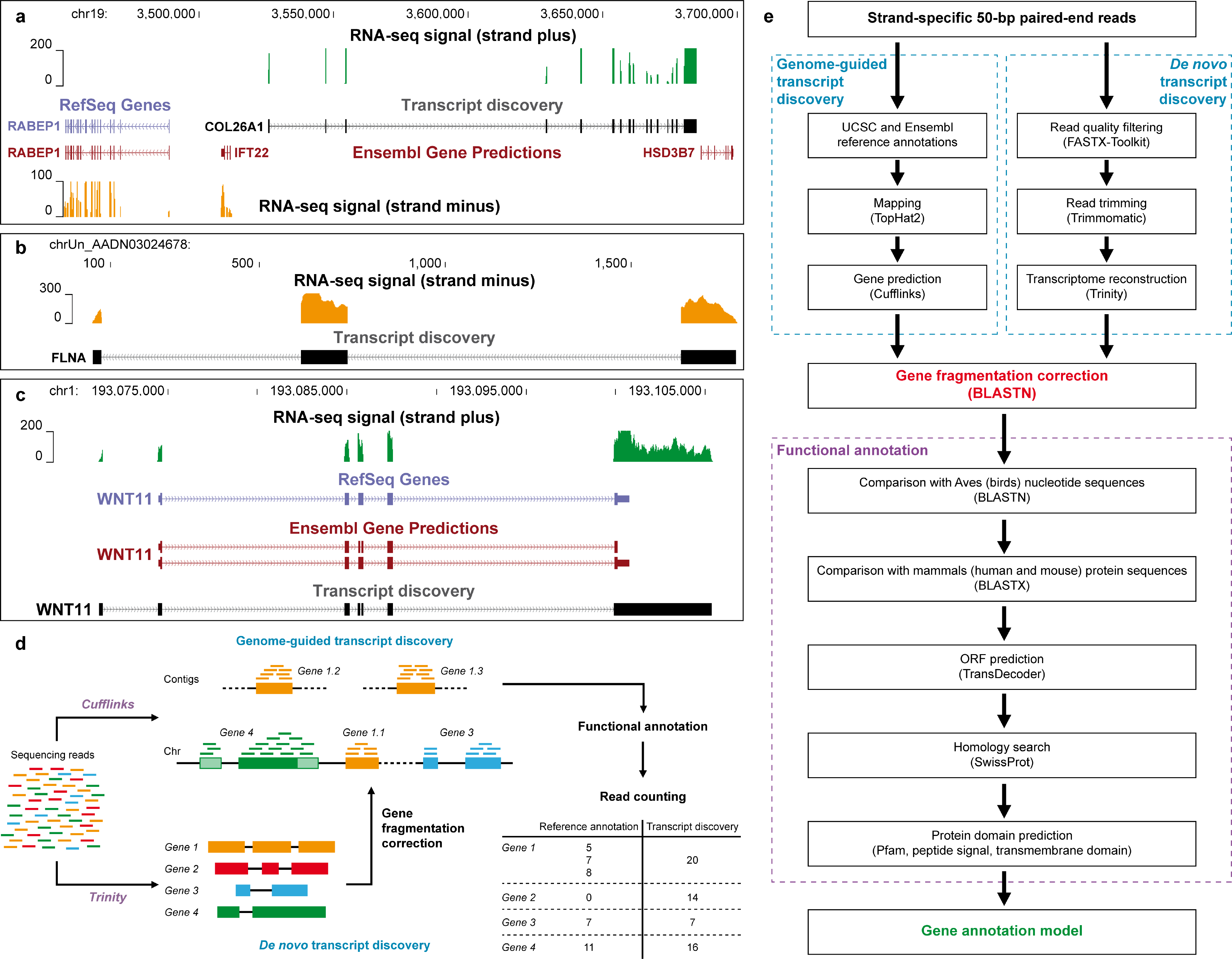
Dual transcript-discovery approach. (**a**) Region surrounding the genes *RABEP1* and *HSD3B7* on chromosome 19. RNA-seq signal on strand plus (green), which does not overlap any gene from UCSC and Ensembl reference annotations, corresponds to the gene *COL26A1*. (**b**) RNA-seq signal (orange) on strand minus of an uncharacterized contig delimitating 3 exons of the gene *FLNA*. (**c**) Region of the gene *WNT11* on chromosome 1. As visible from the RNA-seq signal on strand plus (green), both UCSC and Ensembl reference annotations lack an exon of the 5’-UTR and display a shorter 3’-UTR. (**d**) The dual transcript-discovery approach combined a genome-guided gene prediction with a *de novo* transcriptome reconstruction. This dual approach enabled us to correct for gene fragmentation (orange), to identify missing genes (red) and to adjust existing annotated genes (green), thus improving the assignment rate of RNA-seq read pairs. (**e**) Workflow to design the comprehensive gene annotation model.

In order to improve the read assignment rate, we first performed a genome-guided transcript discovery. This approach was intended to determine more accurately exon-intron junctions, to correct or to complete existing annotated genes, and to identify unannotated genes from the UCSC/Ensembl gene annotation model (Fig. 1d, e). Following this approach, 77.9% of the sequencing read pairs were assigned to gene features, corresponding to 89.8% of the read pairs that were mapped against the genome (Table 1). Therefore, the genome-guided transcript discovery enabled us to raise the read assignment rate by 15.7% as compared to when using both UCSC and Ensembl reference annotations (Table 1). In contrast to genome-guided transcript prediction, *de novo* transcriptome reconstruction relies on overlaps between the sequencing reads to build consensus transcripts, independently of the genome sequence. We therefore applied a genome-independent strategy in combination to the genome-guided approach in order to detect transcripts or regions that were not recovered from the genome sequence, such as those located within gaps or uncharacterized chromosomes (Fig. 1d, e). Reconstructed transcripts thus generated were then compared to the genes obtained with the genome-guided approach in order to remove redundant sequences. Full-length transcripts or transcript regions of at least 400 bp that were not assigned to any gene were extracted and grouped as an artificial chromosome. 4.0% of read pairs were found to map against this additional chromosome and 90.2% of these mapped read pairs were assigned to gene features (Table 1). By considering both transcript-discovery approaches, 90.7% of total read pairs were mapped against the galGal4 chicken genome (86.7%) and reconstructed chromosome (4.0%) (Table 1). 77.9% and 3.6% of read pairs were assigned to gene features from the genome-guided and *de novo* transcript-discovery approaches, respectively (Fig. 2a; Table 1). Therefore, 81.5% of read pairs were assigned to gene features by using this newly established gene annotation model. Given that 62.2% of sequencing read pairs were assigned to gene features by using both UCSC and Ensembl reference annotations, our transcript reconstruction model enabled us to assign 19.3% more read pairs to gene features (Fig. 2a; Table 1).

**Fig. 2.**
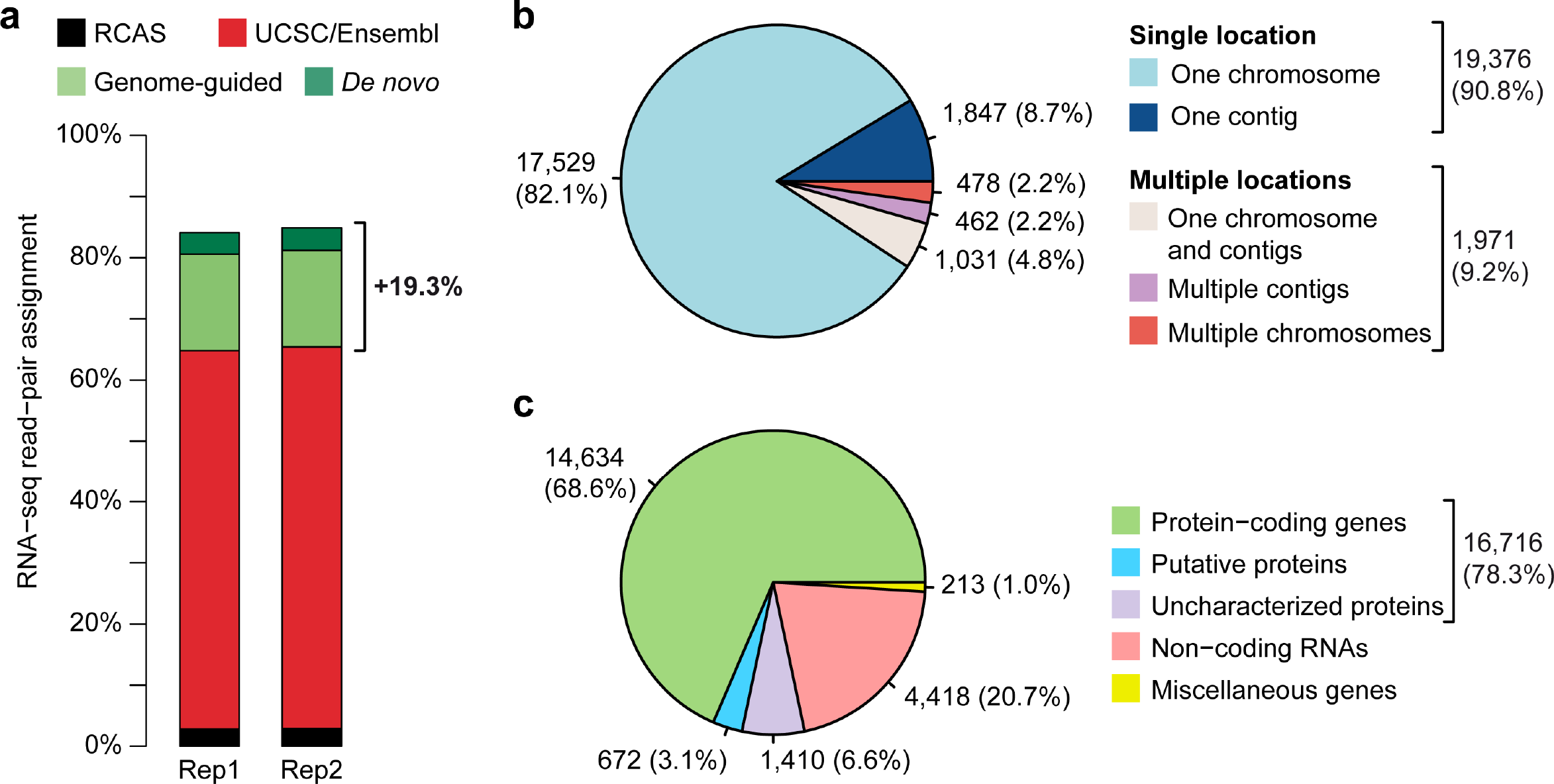
Characteristics of the new gene annotation model. (**a**) The dual transcript-discovery approach combining genome-guided gene prediction (light green) and *de novo* transcriptome reconstruction (dark green) raised the read-pair assignment rate by 19.3% as compared to when using the UCSC and Ensembl reference annotations (red). The proportion of read pairs coming from the RCAS-BP(A) replication competent retroviruses are depicted in black. (**b**) Proportion of gene location on chromosomes and contigs of the chicken reference genome galGal4. 9.2% of identified genes are fragmented due to their location on multiple chromosomes and contigs. (**c**) Proportion of annotated gene biotypes. Most of the annotated genes potentially encode proteins (78.3%). Putative proteins correspond to genes for which at least one protein domain could be detected (3.1%). Uncharacterized proteins are genes with an ORF of at least 100 amino acids without protein domain identified (6.6%). Genes with no sufficient predicted ORF (less than 100 amino acids) are classified as non-coding RNAs (20.7%). Genes encoding spliceosome complex members and ribosomal RNAs, as well as pseudogenes are classified as miscellaneous genes (1.0%).

The genome-independent transcript assembly enabled us to correct for gene fragmentation by gathering gene regions located on multiple chromosomes and contigs together (Fig. 1d, e). In contrast to genome-guided transcript discovery, *de novo* reconstruction of transcripts was not limited by the quality of the reference genome sequence. By comparing transcripts generated from both reconstruction approaches, we were therefore able to group dispersed gene features belonging to a same gene together. Although 19,376 (90.8%) genes were found on one single chromosome or unplaced contig, 1,971 (9.2%) genes were identified as being fragmented (Fig. 2b). These fragmented genes included 478 (2.2%) genes that were located on multiple ordered chromosomes, 462 (2.2%) genes split among multiple unplaced contigs, and 1,031 (4.8%) genes with regions located on one ordered chromosome and additional unplaced contigs (Fig. 2b).

Transcript prediction and reconstruction approaches did not provide any information on gene name and function. Therefore, genes identified by the dual transcript-discovery approach were then annotated by database comparison and protein domain prediction (Fig. 1e). Genes were first compared to bird gene sequences, taking advantage of the recent increase of available genomic data within avian species and their high DNA sequence conservation [20-27]. Undefined genes were then compared at the protein level against mouse and human databases. Finally, prediction of open reading frames (ORFs) and protein domains was performed on remaining unannotated genes by using homology search against SwissProt and Pfam databases, and sequence analysis tools to identify transmembrane domains and peptide signals. Overall, the computed gene annotation model was mostly constituted of protein-coding genes (16,716, 78.3%) (Fig. 2c). However, 672 (3.1%) genes were only partly annotated (putative proteins having at least one protein domain detected), while 1,410 (6.6%) genes remained unannotated (uncharacterized proteins with no protein domain identified but an ORF of at least 100 amino acids). Remaining genes corresponded to miscellaneous genes (213, 1.0%; such as spliceosome complex members, ribosomal RNAs and pseudogenes) and non-coding RNAs (ncRNAs; 4,418, 20.7%) for which no sufficient ORF could be predicted (Fig. 2c). Altogether, this dual transcript-discovery approach enabled us to define an annotation model of 21,347 unique genes. This gene annotation model was composed of 4,029 additional genes as compared to the UCSC and Ensembl reference annotations associated with the galGal4 genome version and enabled us to retrieve 19.3% more information from the RNA-seq data.

## Discussion

The work presented here describes a dual transcript-discovery approach combining genome-guided gene prediction and *de novo* transcriptome reconstruction, which was applied to improve the assignment rate of RNA-seq data obtained from chicken samples. For the first approach, sequencing read pairs are mapped along the genome followed by a genome-dependent transcript discovery, which computes read coverage and exon-intron junctions from gapped alignments, and distance between both reads of each pair. By contrast, the second approach is carried out independently of the reference genome. Sequencing reads are *de novo* assembled by relying on their overlaps to reconstruct full-length transcripts. Genome-guided transcript discovery is more sensitive than *de novo* transcript reconstruction, but requires a reference genome along which RNA-seq reads are mapped for gene prediction [11,14]. Therefore, the choice of the latter method is obvious when no or incomplete genome sequence is available. In the case of the chicken model with its partial and fragmented genome sequence, the choice of a complementary transcript-discovery approach, combining both genome-guided and -independent methods, appears suitable to improve the RNA-seq data quantification and analysis. While the genome-guided approach contributes to correct existing annotated genes and to identify novel genes, the *de novo* transcript reconstruction compensates for gene fragmentation by associating gene parts located on multiple chromosomes or contigs together; and it identifies gene regions or complete genes that do not belong to the genome sequence due to the presence of gaps or uncharacterized fragments. The new annotation model is composed of 21,347 genes, accounting for 4,029 additional genes as compared to the UCSC and Ensembl reference annotations associated with the galGal4 genome version. 1,971 (9.2%) genes have parts spread on multiple locations, while 3,340 (15.6%) genes are identified among the 16,000 unplaced contigs that are not assigned to any ordered chromosome. In addition, the resulting gene annotation model increased the assignment rate of RNA-seq read pairs by 19.3% as compared to when using both galGal4 reference annotations, thus contributing to a more accurate estimation of transcript abundance.

Genome-guided gene prediction and *de novo* transcript reconstruction cannot provide information on either gene name or function, which renders the identification of relevant target genes in downstream analyses difficult. The recent genome sequencing of the zebra finch [25], the turkey [20], the pigeon [24], the falcon [26], the duck [21], and a wide range of additional avian species [22,27] have provided extensive insights into evolutionary and adaptive traits within birds. DNA conservation of protein-coding genes among avian species considerably facilitated the annotation of the 21,347 genes identified by the dual transcript-discovery approach. By combining DNA sequence comparison against avian genes with protein sequence comparison against mammal species and protein domain prediction, 14,847 (69.6%) genes could be assigned and 672 (3.1%) putative protein-coding genes could be identified. The 5,828 (27.3%) remaining genes were divided between uncharacterized proteins and ncRNAs depending on the length of the predicted ORF. However, genes encoding uncharacterized proteins could be also potentially non-coding since none of the protein domains investigated was detected within their putative ORF. On the other hand, ncRNAs remain challenging to annotate according to a recent study comparing an extensive repertoire of long multi-exonic ncRNAs across 11 tetrapods separated by up to 370 million years [28]. Besides their overall weak conservation as compared to protein-coding sequences, long ncRNAs display high tissue specificity and rapidly diverge through evolution, which renders their annotation difficult by comparing with other species.

Since the first draft released in 2004, the International Chicken Genome Consortium has improved the *Gallus gallus* reference genome [1]. In December 2015, the fifth version of the chicken genome (galGal5) was released [29]. As compared to the fourth version, this release is 200 Mb longer and includes 3 additional chromosomes (30, 31, 33) but remains highly fragmented. Indeed, this fifth version is still composed of 15,400 unassigned contigs and 8,000 contigs assigned with low confidence, accounting for about 17% of the total genome size. Improvement of the chicken genome is an on-going project and a new version should be released within the next few years. It is reasonable to believe that continuing efforts will contribute to elucidate the full sequence of the chicken genome in a near future. Until then, applying the dual transcript-discovery approach described here prior to the analysis of RNA-seq data *per se* enhances the sensitivity of gene expression profiles. This is particularly relevant considering that genes and splicing variants are specifically expressed in certain cell types or tissues, at different developmental stages and conditions within a single organism. For instance, we used the gene annotation model presented here as guide in a recent study, where we aimed at identifying genes that were regulated upon overexpression of connective tissue-associated transcription factors in chick micromass cultures (Orgeur *et al.,* in preparation). More broadly, this approach could be also employed to analyse RNA-seq data of other organisms lacking manually-curated, high-quality reference annotation.

## Methods

### Chick embryos

Fertilized chick eggs were obtained from VALO BioMedia (Lohmann Selected Leghorn strain, Osterholz-Scharmbeck, Germany). Chick embryos were staged according to the number of days *in ovo* at 37.5°C.

### Chick micromass cultures

Two independent biological replicates of micromass cultures were prepared from limb buds of E4.5 chick embryos, infected with RCAS-BP(A) retroviruses carrying no recombinant protein and cultivated for 5 days as described previously [30,31]. Briefly, ectoderm was dissociated by using a Dispase solution (Gibco) at 3 mg/mL and limb mesenchyme was digested by using a solution composed of 0.1% Collagenase type Ia (Sigma-Aldrich), 0.1% Trypsin (Gibco) and 5% FBS (Biochrom) in DPBS (Gibco). Prior to seeding, mesenchymal cells were mixed with retroviruses and maintained in culture for 5 days at 37°C in DMEM/Ham’s F-12 (1:1) medium (Biochrom) supplemented with 10% FBS, 0.2% chicken serum (Sigma-Aldrich), 1% L-glutamine (Lonza) and 1% penicillin/streptomycin (Lonza).

### RNA sequencing

For both replicates, RNA extracts were obtained by harvesting 6 micromass cultures with RLT buffer (Qiagen). Total RNAs were purified by using the RNeasy mini kit (Qiagen) in combination to a DNase I (Qiagen) treatment to prevent genomic DNA contamination. RNA libraries were prepared by using the TruSeq Stranded mRNA Library Preparation kit (Illumina), which enables to preserve the RNA strand orientation. Strand-specific 50-bp paired-end reads were generated by using a HiSeq 2500 sequencer (Illumina) with a mean insert size of 150 bp.

### Genome-guided transcript discovery

RNA-seq data obtained from both biological replicates of micromass cultures were processed independently. Strand-specific read pairs were mapped against the chicken genome galGal4 [1] by using TopHat2 v0.14 [32] (parameters: −r 150; −N 3; −−read-edit-dist 3; −−library-type fr-firststrand; −i 50; −G). UCSC (galGal4) and Ensembl (release 75) annotations were downloaded from Illumina iGenomes [33] and compared by using Cuffcompare from the Cufflinks suite v2.1.1 [8]. Identical genes were retrieved only once and merged with the unique genes from each annotation. In case of discordant genes, the gene annotation with the best coverage was selected. The resulting gene annotation model composed of 17,318 genes was used as input for TopHat2 mapping. Transcript discovery was performed for each replicate by using Cufflinks v2.1.1 [8] (parameters: −b; −u; −library-type, fr-firststrand; −g) and the combined gene annotation model as guide. Resulting annotations were merged into a single model by using the Cufflinks tool Cuffmerge v2.1.1 [8] (Additional file 1).

### *De novo* transcript discovery

A second transcript-discovery approach was led independently of the genome sequence. Low-quality RNA-seq reads from each replicate of micromass cultures were first filtered out by using the FASTX-Toolkit v0.0.13 [34]. Reads with a median quality value lower than 28 were discarded. Filtered read pairs were then trimmed by using Trimmomatic v0.32 [35] (parameters: ILLUMINACLIP TruSeq3 paired-end for HiSeq, seedMismatches 2, palindromeClipThreshold 30, simpleClipThreshold 10; LEADING 5; TRAILING 5; MINLEN 36). Complete read pairs were then assembled by using Trinity r20140717 [16] (default parameters except for the strand-specific library orientation set at RF). Resulting contigs were compared to the gene sequences obtained by the first approach by using BLASTN from BLAST+ v2.2.31+ [36] (parameters: −strand plus; −dust no; −soft_masking no). Contigs were assigned to a given gene if they matched at least 40 bp with a percentage of identities higher than 90%. Assigned contigs that were not fully covered by a given gene were further processed to extract continuous uncovered regions of at least 400 bp. Remaining contigs were mapped against the galGal4 genome by using BLASTN. Contigs were assigned to a given gene if they were located between two gene features, potentially corresponding to an exon missed by Cufflinks, or in the vicinity of a first or last exon, potentially corresponding to a missing 5’- or 3’-untranslated region (UTR), respectively. Remaining unmapped contigs were retrieved as they could correspond to non-defined genomic regions. Unmapped, unassigned and non-covered contigs or regions were compared to each other to remove redundant sequences. Unique contig sequences were gathered together as an artificial chromosome and separated to each other by 250 bp of nucleotides N, corresponding to the total length of read pairs (50 bp for each read and 150 bp as insert size) (Additional files 2 and 3).

### Functional annotation

Gene sequences retrieved from both transcript-discovery approaches were then compared to existing databases for gene name assignment. First, genes were compared to the NCBI RefSeq transcript database by using BLASTN (parameters: −strand plus; −dust no; −soft_masking no). Comparison was limited to Aves (birds) sequences (taxid 8782). Genes with a percentage of identities higher than 90% for chicken genes or 75% for bird genes, and bidirectionaly covered on at least 50% of their length were assigned to the corresponding hits. Non-annotated gene sequences were then compared against the NCBI human (taxid 9606) and mouse (taxid 10090) non-redundant protein database by using BLASTX from BLAST+ v2.2.31+ [36] (parameters: −strand, plus; −seg, no). Genes with a percentage of homology of at least 30% and covered by at least 50% of their length were filtered. Matching protein accession numbers were converted into gene accession numbers by using the Hyperlink Management System [37]. ORF prediction was finally performed on remaining genes by using TransDecoder v2.1.0 [38] (strand specificity parameter: −S). ORFs of at least 100 amino acids were annotated by using Trinotate v3.0.1 [39]. Functional annotation was based on the following protein predictions: (i) BLASTX and BLASTP homology search against the SwissProt database [40]; (ii) protein domain prediction against the Pfam database [41] by using HMMER v3.1b2 [42]; (iii) peptide signal prediction by using SignalP v4.1 [43]; and (iv) transmembrane domain prediction by using tmHMM v2.0c [44]. Resulting functional annotation was divided into three categories: (i) putative proteins, for which at least one protein domain could be identified; (ii) uncharacterized proteins, corresponding to ORFs for which no protein domain could be identified; and (iii) ncRNAs, corresponding to genes with an ORF shorter than 100 amino acids.

### Fragment counting

Strand-specific read pairs mapped against the galGal4 genome and the artificial chromosome generated from the *de novo* transcript discovery were first split by strand by using SAMtools v1.2 [45] according to their FLAG field (strand plus: −f 128 −F 16, −f 80; strand minus: −f 144, −f 64 −F 16). Fragments (both reads of a pair) mapped on gene features were counted by using featureCounts v1.4.6-p3 [46] (parameters: −p; −s 2; −−ignoreDup; −B; −R). Chimeric fragments aligned on different chromosomes were taken into consideration to overcome the gene fragmentation due to the location of gene parts on multiple chromosome contigs.

### Additional files

**Additional file 1:** gene annotation model in GTF format associated with the galGal4 version of the chicken genome.

**Additional file 2:** artificial chromosome in FASTA format containing the unique contig sequences generated from the *de novo* transcript discovery. Unique contig sequences are separated to each other by 250 bp.

**Additional file 3:** gene annotation model in GTF format associated with the artificial chromosome generated from the *de novo* transcript discovery.

## Acknowledgements

We thank Georgeta Leonte (Freie Universität, Berlin, Germany) for helping preparing and collecting the samples, as well as Stéphane Descorps-Declère and Marc Monot (Institut Pasteur, Paris, France) for critical reading of the manuscript. We are grateful to the Sequencing Core Facility of the Max Planck Institute for Molecular Genetics for processing the RNA-seq. We are thankful to Peter Hansen and Peter N. Robinson (Charité Universitätsmedizin, BCRT, Berlin, Germany), as well as Marius van den Beek and Christophe Antoniewski (Institut de Biologie Paris-Seine, ARTbio, Paris, France) for providing access to Galaxy web servers.

## Availability of data

Sequencing data have been deposited on the Gene Expression Omnibus (GEO) database under the SuperSeries accession number GSE100517. Both samples that have been used for this study are available under the SubSeries GSE100516 via the accession numbers GSM2685833 and GSM2685834.

## Authors’ contributions

MO, DD and SS designed and conceived the study. MO performed the experiments and collected the samples. MO and MM analysed the data. STB and BT performed and supervised the RNA-seq procedure. MO, DD and SS wrote the manuscript, with comments and approval from all authors.

